# Engineering a 3D hydrogel system to study optic nerve head astrocyte morphology and behavior in response to glaucomatous insult

**DOI:** 10.1101/2022.01.08.474536

**Authors:** Ana N. Strat, Alexander Kirschner, Hannah Yoo, Ayushi Singh, Tyler Bagué, Haiyan Li, Samuel Herberg, Preethi S. Ganapathy

**Affiliations:** Department of Ophthalmology & Visual Sciences, SUNY Upstate Medical University, Syracuse, NY 13210, USA; Department of Neuroscience and Physiology, SUNY Upstate Medical University, Syracuse, NY 13210, USA; BioInspired Institute, Syracuse University, Syracuse, NY 13244, USA; Department of Cell and Developmental Biology, SUNY Upstate Medical University, Syracuse, NY 13210, USA; Department of Biochemistry and Molecular Biology, SUNY Upstate Medical University, Syracuse, NY 13210, USA; Department of Biomedical and Chemical Engineering, Syracuse University, Syracuse, NY 13244, USA

**Keywords:** reactive gliosis, transforming growth factor beta 2, extracellular matrix, GFAP, fibronectin, collagen IV

## Abstract

In glaucoma, astrocytes within the optic nerve head (ONH) rearrange their actin cytoskeleton, while becoming reactive and upregulating intermediate filament glial fibrillary acidic protein (GFAP). Increased transforming growth factor beta 2 (TGFβ2) levels have been implicated in glaucomatous ONH dysfunction. A key limitation of using conventional 2D culture to study ONH astrocyte behavior is the inability to faithfully replicate the in vivo ONH microenvironment. Here, we engineer a 3D ONH astrocyte hydrogel to better mimic in vivo mouse ONH astrocyte (MONHA) morphology, and test induction of MONHA reactivity using TGFβ2. Primary MONHAs were isolated from C57BL/6J mice and cell purity confirmed. To engineer 3D cell-laden hydrogels, MONHAs were mixed with photoactive extracellular matrix components (collagen type I, hyaluronic acid) and crosslinked for 5 minutes using a photoinitiator (0.025% riboflavin) and UV light (405-500 nm, 10.3 mW/cm2). MONHA-encapsulated hydrogels were cultured for 3 weeks, and then treated with TGFβ2 (2.5, 5.0 or 10 ng/ml) for 7 days to assess for reactivity. Following encapsulation, MONHA retained high cell viability in hydrogels and continued to proliferate over 4 weeks as determined by live/dead staining and MTS assays. Sholl analysis demonstrated that MONHAs within hydrogels developed increasing process complexity with longer process length over time. Cell processes connected with neighboring cells, coinciding with Connexin43 expression within astrocytic processes. Treatment with TGFβ2 induced reactivity in MONHA-encapsulated hydrogels as determined by altered F-actin cytoskeletal morphology, increased GFAP expression, and elevated fibronectin and collagen IV deposition. Our data sets the stage for future use of this 3D biomimetic ONHA-encapsulated hydrogel to investigate ONHA behavior in response to glaucomatous insult.

## 1. Introduction

Glaucoma is a chronic progressive optic neuropathy that leads to irreversible blindness due to the loss of retinal ganglion cells (RGCs) (Neuman et al., 2014; Weinreb et al., 2014). The main risk factor for the disease is elevated intraocular pressure (IOP). As such, current medical and surgical interventions are designed to lower IOP to prevent disease progression (Stein et al., 2021; Weinreb et al., 2014). Much of the mechanism of how IOP affects RGC viability remains unknown. RGCs are initially damaged within the optic nerve head (ONH) (Crawford Downs et al., 2011; Neuman et al., 2014; Weinreb et al., 2014), which contains lamina cribrosa cells, astrocytes, microglia, and an extracellular matrix (ECM) network that supports RGC axons as they leave the globe through the optic nerve (Hernandez et al. 1986; Hernandez et al. 1988). IOP elevation in glaucoma strongly correlates with aberrant ONH ECM remodeling and increased mechanical stress on RGC axons. Of the cells within the ONH, astrocytes are likely candidates for transducing IOP insult into changes in ECM structure. Moreover, ONH astrocytes have been identified as key modulators of RGC axonal health in both early and late stages of disease (Clarke et al., 2018, Cooper et al., 2018).

Early in glaucoma, it is likely that astrocytes are protective to RGCs (Sun et al., 2017; Cooper et al., 2020). Immediately after IOP elevation, astrocytes within the ONH become reactive and upregulate intermediate filament glial fibrillary acidic protein (GFAP) levels (Cooper et al., 2020). Astrocytes additionally rearrange their actin cytoskeleton and cellular processes, coinciding with Connexin 43 (CX43) gap-junction coupling, to promote axonal health (Cooper et al., 2018; Sun et al., 2017; Tehrani et al., 2016; Tehrani et al., 2019). However, later in the disease, there is evidence that excessive astrocytic reactivity is neurotoxic to RGCs (Sloan and Barres, 2014; Liddelow et al., 2017; Sterling et al., 2020). Reactive astrocytes secrete both ECM crosslinking and degrading enzymes and upregulate pro-fibrotic cytokines such as transforming growth factor beta 2 (TGFβ2), which can significantly impact ECM integrity (Hernandez et al., 2000). Many studies implicate a role of elevated TGFβ2 in ECM remodeling within the glaucomatous ONH (Kim et al., 2017; Pena, et al. 1999; Zode et al., 2011). Thus, careful investigation of the relationship between glaucomatous insult, ONH astrocyte behavior, and the surrounding ECM is integral to understanding glaucoma pathobiology. Our overarching goal is to develop a model system ideal for investigating this relationship.

Monolayer ONH astrocyte cultures subjected to biochemical (i.e., TGFβ2 treatment) or mechanical (i.e., hydrostatic pressure) stressors frequently display increased reactivity, actin remodeling, and upregulation of ECM protein production (Ricard et al., 2000). However, there are significant limitations to studying glaucoma pathophysiology using 2D monolayer cultures. Astrocytes, when cultured on supraphysiologically stiff substrates such as glass or plastic (Caliari and Burdick, 2016) display a reactive phenotype and lack their characteristic stellate morphology. Additionally, they fail to form typical star shape in 2D (Hernandez et al., 2000). In an attempt to better mimic *in vivo* astrocyte stellate morphology and to limit baseline reactivity, a few groups have used viscoelastic hydrogel systems (i.e., water-swollen networks of natural or synthetic polymers) to provide more biologically relevant substrates for cell growth (Placone et al., 2015; Mulvihill et al., 2017). Few preliminary studies have described rat optic nerve head astrocyte-encapsulated hydrogels (Boazak et al., 2019; Foltz et al., 2021). Our work seeks to further the existing literature by encapsulating mouse ONH astrocytes inside the 3D polymer network, systematically characterizing time-dependent process complexity, and analyzing cellular reactivity in response to glaucomatous stressors.

We recently described a novel tissue-engineered 3D trabecular meshwork hydrogel system (Li et al., 2021) by mixing human trabecular meshwork cells with ECM biopolymers (collagen type I, hyaluronic acid; HA, and elastin-like polypeptide) followed by photoinitiator-mediated short UV crosslinking. Building on this previous work, in the present study we engineer a hydrogel-based model system containing mouse ONH astrocytes (MONHAs) and photoactive ECM proteins (collagen type I and HA) that is crosslinked with UV light using 0.025% riboflavin as photoinitiator to (1) better mimic ONH astrocyte stellate morphology and cell-cell interactions, and (2) reliably induce ONH astrocyte reactivity and ECM production in response to relevant glaucomatous stressors. Focusing on astrocyte morphology, we characterized length and complexity of cellular processes over time using Sholl analysis. We investigated whether MONHAs retain astrocyte markers GFAP and CX43 expression after four weeks in culture. Lastly, to test whether encapsulated MONHAs respond to a known glaucomatous stressor, cell-encapsulated hydrogels were treated with TGFβ2 followed by assessment of GFAP expression, actin cytoskeletal remodeling, and ECM protein deposition in 3D.

## 2. Methods

### 2.1 MONHA isolation and culture

C57BL/6J mice were purchased from the Jackson Laboratory (Bar Harbor, ME) and bred in house according to the institutional guidelines for the humane treatment of animals (IACUC #473) and to the ARVO Statement for the Use of Animals in Ophthalmic and Vision Research. For each harvest in this study 6-8 mice aged 6-8 weeks were used. The isolation of primary MONHAs and cell culture was performed as previously described (Kirschner et al. 2021). Briefly, using a SMZ1270 stereomicroscope (Nikon Instruments, Melville, NY), ONH tissue was dissected from each ocular globe proximal to the sclera. Tissue samples were digested in 0.25% trypsin (Invitrogen, 25200-056, Carlsbad, CA) for 15 minutes at 37 °C and then resuspended in MONHA growth medium (Dulbecco’s modified Eagle’s medium, DMEM/F12 (Invitrogen, 11330-032) + 10% fetal bovine serum (Atlanta Biologicals, S11550, Atlanta, GA) + 1% penicillin/streptomycin (Corning, 30-001-CI, Manassas, VA) + 1% Glutamax (Invitrogen, 35050-061, Grand Island, NY) + 25 ng/ml epidermal growth factor (EGF; Sigma, E4127-5X, St. Louis, MO). After digestion ONH tissue was plated on 0.2% gelatin (Sigma, G1393) coated T75 cell culture flasks and kept at 37 °C in a humidified atmosphere with 5% CO_2_. MONHAs migrated from ONH tissue over 10-14 days before first passage. We used passages 2-5 of six separate preparations (MONHA04, MONHA05, MONHA06, MONHA07, MONHA08, MONHA09) for all experiments.

### 2.2 MONHA cell characterization

MONHAs were seeded at 1 × 10^4^ cells/cm^2^ on sterile glass coverslips in 24-well culture plates (Thermo Fisher Scientific, Waltham, MA). After 48 h, cells were fixed with 4% paraformaldehyde (PFA; J19943-K2, Thermo Fisher Scientific) at room temperature for 10 min, and permeabilized with 0.5% Triton X-100 (Thermo Fisher Scientific, 85111) at room temperature for 30 minutes. Cells were washed in Dulbecco’s Phosphate Buffered Saline 1X (DPBS; Invitrogen, 14190-44) and blocked (PowerBlock; Biogenx, HK085-5K, San Ramon, CA) for 1 h at room temperature. Cells were then incubated for 1 hour at room temperature with rabbit anti-glial fibrillary acidic protein (GFAP; rabbit anti-GFAP antibody, 1:300, Dako, Z0334, Carpinteria, CA), rabbit anti-oligodendrocyte specific protein (OSP; rabbit anti-OSP antibody, 1:100, Abcam, Ab53041, Cambridge, MA) or rat anti-F4/80 (rat anti-F4/80 antibody, 1:50, BioRad, MCA497GA, Hercules, CA). Cells were again washed in DPBS and incubated for 1 h at room temperature with Alexa Fluor® 488-conjugated secondary antibody (goat polyclonal antibody to rabbit IgG, 1:500, Abcam, Ab15077). Nuclei were counterstained with 4’,6’-diamidino-2-phenyliondole (DAPI; Invitrogen, D1306). Coverslips were mounted with ProLong™ Gold Antifade (Thermo Fisher Scientific, P36930) on Superfrost™ Plus microscope slides (Fisher Scientific) and fluorescent images were acquired with Eclipse N*i* microscope (Nikon). Four fields of view at 20x magnification were taken from each coverslip per culture. Number of GFAP-, OSP-, or F4/80-positive cells versus total number of cells were quantified from acquired images.

### 2.3 Hydrogel precursor solutions

Methacrylate-conjugated bovine collagen type I (MA-COL; molecular weight: ∼300 kDa, degree of methacrylation: ∼60–70%; Advanced BioMatrix, Carlsbad, CA, USA) was reconstituted according to the manufacturer’s instructions with sterile 20 mM acetic acid at 4 mg/ml. 1 ml MA-COL was neutralized with 90 μl neutralization buffer (Advanced BioMatrix) for hydrogel use. Thiol-conjugated hyaluronic acid (SH-HA; Glycosil®; molecular weight: ∼300 kDa, degree of thiolation: ∼20–30%; Advanced BioMatrix) was reconstituted in sterile diH2O at 10 mg/ml according to the manufacturer’s instructions.

### 2.4 Preparation of PDMS molds

A 10:1 ratio of elastomer to curing agent was prepared according to the manufacturer’s protocol for polydimethylsiloxane (PDMS; Sylgard 184, Dow Corning, Midland, MI, USA). Using a 3D printer (F170; Stratasys, Ededn Prairie, MN, USA), 10 mm diameter x 1 mm depth negative molds were made from ABS-M30 filament. The PDMS mixture was poured into the negative molds and degassed in a desiccator under vacuum before curing overnight at 60 °C. PDMS molds were sterilized prior to use in culture.

### 2.5 Preparation of MONHA-encapsulated hydrogels

MONHAs (2.5 × 10^6^ cells/ml or 5.0 × 10^6^ cells/ml) in media were mixed with 3.1 mg/ml methacrylate-conjugated bovine collagen type I, 1 mg/ml thiol-conjugated hyaluronic acid, and 0.025% (w/v) riboflavin (Sigma, R7774) (photoinitiator) on ice (Fig. 1). The chilled MONHA hydrogel precursor solution was pipetted as (1) 10 μl droplets onto PDMS-coated (Sylgard 184; Dow Corning) 24-well culture plates, (2) 20 μl droplets onto 12 mm round glass coverslips sandwiched with Surfasil (Fisher Scientific) coated coverslips on top, or 80 μl into custom 10 × 1 mm PDMS molds. Constructs were then crosslinked by exposure to UV light (OmniCure S1500 UV Spot Curing System; Excelitas Technologies, Mississauga, Ontario, Canada) at 405-500 nm, 10.3 mW/cm^2^ for 5 minutes. MONHA growth medium was added to each well and replenished every 2-3 days, and constructs were cultured for 1-4 weeks.

**Fig. 1.**
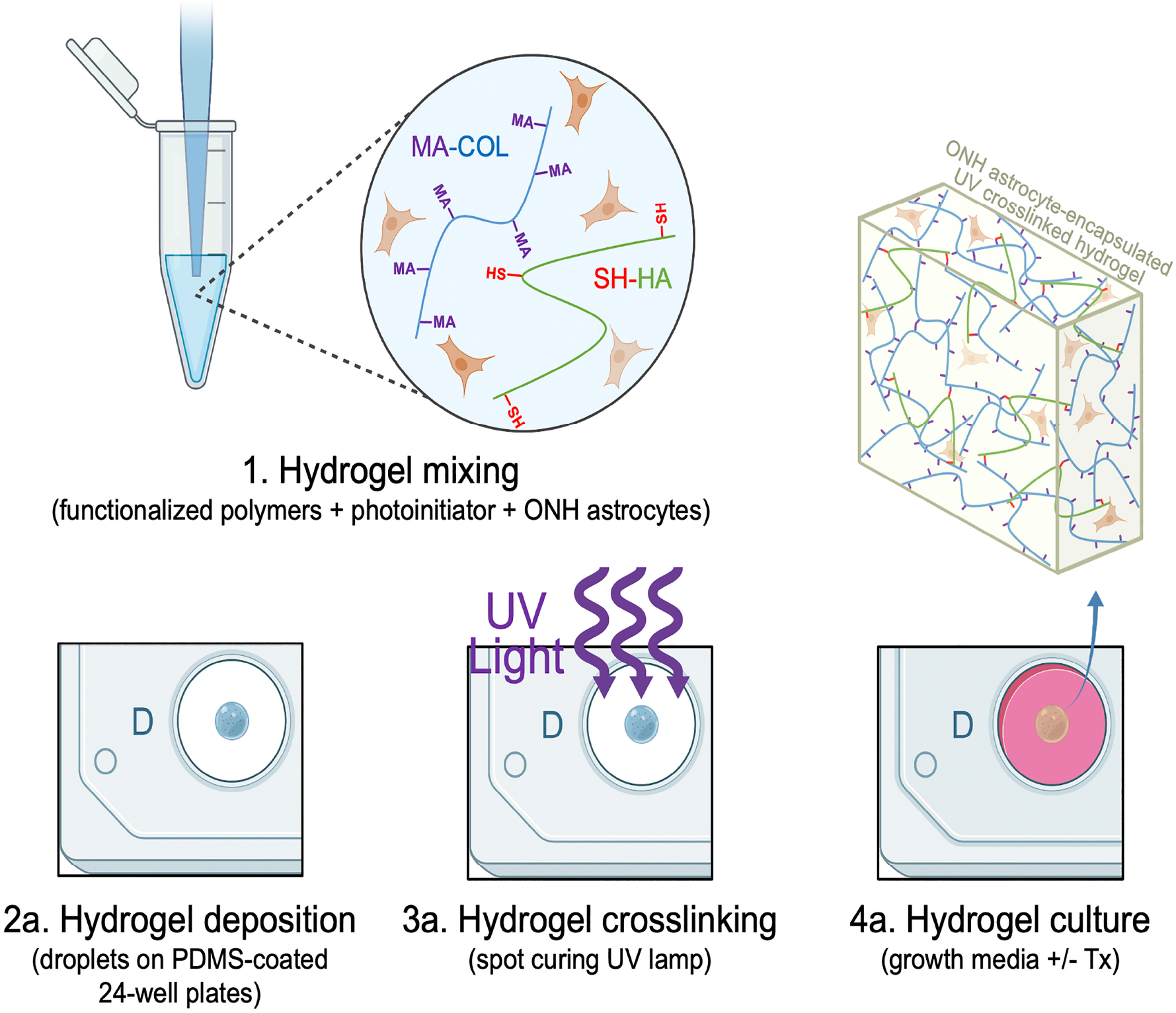
Schematic of MONHA-encapsulated hydrogel formulation.

### 2.6 MONHA hydrogel rheology analysis

Acellular and MONHA encapsulated hydrogels were created using 80 μl of hydrogel mixture per 10 × 1 mm PDMS mold. Samples were UV crosslinked as described in Methods 2.4. Hydrogel viscoelasticity of acellular and MONHA-encapsulated hydrogels (N = 4 per group) was obtained at day 0 using a Kinexus rheometer (Malvern Panalytical, Westborough, MA, USA) fitted with an 8 mm diameter parallel plate. Rheometry measures were performed similarly to Li et al. (2021). Briefly, the 8 mm geometry was lowered on top of the hydrogels to a calibration normal force of 0.02 N, and an oscillatory shear-strain sweep test (0.1–60%, 1.0 Hz, 25 °C) determined values for storage modulus (G’) and loss modulus (G”) in the linear region. Storage modulus for each sample was then converted to elastic modulus (E) calculated as E = 2 * (1 + v) * G′, with Poisson’s ratio (v) of 0.5 for the ECM hydrogels assumed (Lodge et al., 2020; Li et al., 2021(b)).

### 2.7 MONHA encapsulated hydrogel cell viability and proliferation analysis

Using a LIVE/DEAD™ Viability/Cytotoxicity Kit (i.e., live = green-stained, dead = red-stained) (Invitrogen), cell viability was assessed. MONHA hydrogels were incubated at 37 °C for 45 min with the staining solutions (calcein-AM (0.5 μl/ml) and ethidium homodimer-1 (2 μl/ml) diluted in media according to the manufacturer’s instructions and then washed with DPBS. Fluorescent images were captured after initial UV crosslinking on day 0 (N = 3 per group), and weeks 1-4 with an Eclipse *Ti* microscope (Nikon). Four quadrants per hydrogel were imaged and analyzed on day 0 to quantify percent MONHA cell viability (i.e., ratio of live to total cells). Cell proliferation was quantified with the CellTiter 96® Aqueous Non-Radioactive Cell Proliferation Assay (Promega, Madison, WI, USA) as per manufacturer’s instructions. MONHA encapsulated hydrogels were incubated with the staining solution (38 μl MTS, 2 μl PMS solution, 200 μl media) at 37 °C for 90 minutes. Absorbance at 490 nm was then assessed with a spectrophotometer plate reader (BioTEK, Winooski, VT, USA). Fold-change over time of blank-corrected absorbance values was analyzed to quantify cell proliferation in hydrogels.

### 2.8 MONHA hydrogel cell morphology and immunocytochemistry analysis

MONHA-encapsulated hydrogels on glass coverslips were fixed with 4% PFA at 4 °C overnight, permeabilized with 0.5% Triton™ X-100 (Thermo Fisher Scientific), blocked with blocking buffer (BioGeneX), and stained for filamentous F-actin, GFAP, or CX43 as previously described (Li et al., 2021; Kirschner et al., 2021). Briefly, MONHA-encapsulated hydrogels were stained for F-actin with Alexa fluor® 488-or 594-conjugated Phalloidin (1:500, Abcam, Ab176757 or Ab176753) and primary antibody against GFAP (rabbit anti-GFAP, 1:100, Dako, Z0334, Carpinteria, CA) or CX43 (rabbit anti-CX43, 1:100, Cell Signaling Technologies, 3512) overnight, followed by incubation with an Alexa Fluor® 488-conjugated secondary antibody (goat polyclonal antibody to rabbit IgG, 1:500, Abcam, Ab150077). Nuclei were counterstained with DAPI, and fluorescent images were acquired using an Eclipse N*i* microscope (Nikon).

### 2.9 Confocal Microscopy and 3D Analysis

Images of MONHA-encapsulated hydrogels were captured using Zeiss LSM510 scanning confocal microscope. The image size was set to 1024 × 1024 pixels in x/y with a resolution of 0.42 μm per pixel. Individual z-stacks consisted of 5 slices with the z-step interval set to 1.5 μm. The analysis for signal intensity was determined using Z-project Maximum Intensity Projection in Fiji (NIH, Bethesda, MD) across individual z-stacks.

### 2.10 MONHA-encapsulated hydrogel cell morphology analysis

Phalloidin-stained confocal z-stack images were analyzed in Fiji. Tracings of cell processes and branching were performed using Fiji plugin NeuronJ. Sholl analysis was conducted to evaluate process complexity, i.e., the number of astrocytic processes intersecting concentric spheres originating from the cell body (Lye-Barthel et al., 2013). Total mean process length (mean of total processes + branches) and degree of branching (number of primary processes and branches/ number of primary processes) were analyzed for each cell (Placone et al., 2015).

### 2.11 MONHA-encapsulated hydrogel treatments and analysis of F-actin levels

MONHA-encapsulated hydrogels were cultured in MONHA growth medium for 21 days followed by treatment with increasing doses of TGFβ2 (vehicle control, 2.5 ng/ml, 5 ng/ml, and 10 ng/ml; R&D Systems, Minneapolis, MN) for 7 days. Hydrogels were subsequently stained with Alexa fluor® 488-or 594-conjugated Phalloidin and confocal imaging acquired as described in Methods 2.8. Fold changes in normalized F-actin expression were analyzed from quantification over four fields of view per coverslip with background subtraction.

### 2.12 MONHA-encapsulated hydrogel sectioning and immunohistochemical analysis

MONHA-encapsulated hydrogels cultured for 21 days were treated with TGFβ2 for 7 days before 4% PFA fixation at 4 °C overnight. Subsequently, hydrogels were incubated in 30% sucrose at 4 °C for an additional 24 hours. They were then washed with DPBS and embedded in Tissue-Plus™ O.C.T. Compound (Fisher Scientific) before flash freezing in liquid nitrogen. Using a cryostat (Leica Biosystems Inc., Buffalo Grove, IL, USA) 20 μm sections were cut and collected on Superfrost™ Plus microscope slides (Fisher Scientific). Cellular hydrogel sections were permeabilized with 0.5% Triton™ X-100, incubated in blocking buffer for 1 hour, and then immunostained for either fibronectin (rabbit anti-fibronectin antibody, 1:500, Abcam, Ab45688), collagen IV (rabbit anti-collagen IV, 1:500; Abcam, Ab6586), or GFAP (rabbit anti-GFAP, 1:100, Dako) overnight. Sections were then stained with Alexa Fluor® 488-conjugated secondary antibody (goat polyclonal antibody to rabbit IgG, 1:500, Abcam, Ab150077) for 1 hour; nuclei were counterstained with DAPI. Slides were mounted with ProLong™ Gold Antifade (Thermo Fisher Scientific), and fluorescent images were acquired using an Eclipse Ni microscope (Nikon) or Zeiss LSM510 scanning confocal microscope. Fold changes in normalized signal intensity were analyzed from quantification over three fields of view per section with background subtraction using FIJI (NIH, Bethesda, MD), as described in Methods 2.9.

### 2.13 Statistical analysis

Individual sample sizes are specified in each figure caption. Comparisons between groups were assessed by unpaired t-test, one-way, two-way, or main effect analysis of variance (ANOVA) with Tukey’s multiple comparisons *post hoc* as appropriate. A two-way ANOVA main effect only model was used to analyze process complexity over time (i.e., 28 d) (Alexander et al., 2016). All data are shown with mean ± SD. The level of significance was set to p < 0.05 or lower. GraphPad Prism software v9.2 (GraphPad Software, La Jolla, CA, USA) was used for all analyses.

## 3. Results

### 3.1 MONHA-encapsulated hydrogel stiffness is within the range of neurological tissues

Primary astrocytes were isolated and cultured from ONH tissue from 6-8 weeks old C57BL/6J mice, and cell purity was confirmed as previously described (Suppl. Fig. 1) (Kirschner et al., 2021). Thusly characterized MONHAs were used for all subsequent hydrogel experiments. Astrocytes are abundantly present within the ONH, and they form a glial lamina occupying roughly half of the lamina cribrosa volume (Sun et al, 2009; Vecino et al., 2016). To determine optimal density of astrocytes to encourage stellate morphology and cell-cell interaction, we encapsulated 2.5 × 10^6^ cells/ml and, separately, 5 × 10^6^ cells/ml within our hydrogels. The tissue stiffness within central nervous system and ONH tissues range from 0.1 – 1.4 kPa (Budday et al., 2015; Budday et al., 2017). Previous astrocyte-encapsulated viscoelastic hydrogels have reported stiffnesses ranging from 0.0423 to 0.991 kPa (Hu et al., 2021). To determine the stiffness of our hydrogel system, we performed rheology measures on both acellular and MONHA encapsulated hydrogels. Immediately after UV crosslinking, acellular hydrogels had an elastic modulus of 0.201 – 0.319 kPa, while 2.5 × 10^6^ cells/ml MONHA encapsulated hydrogels had an elastic modulus range of 0.310 – 0.368 kPa (Fig. 2A). Thus, MONHA-encapsulated hydrogels harbored stiffnesses within the physiologic range of neurologic and ONH tissues.

**Fig. 2.**
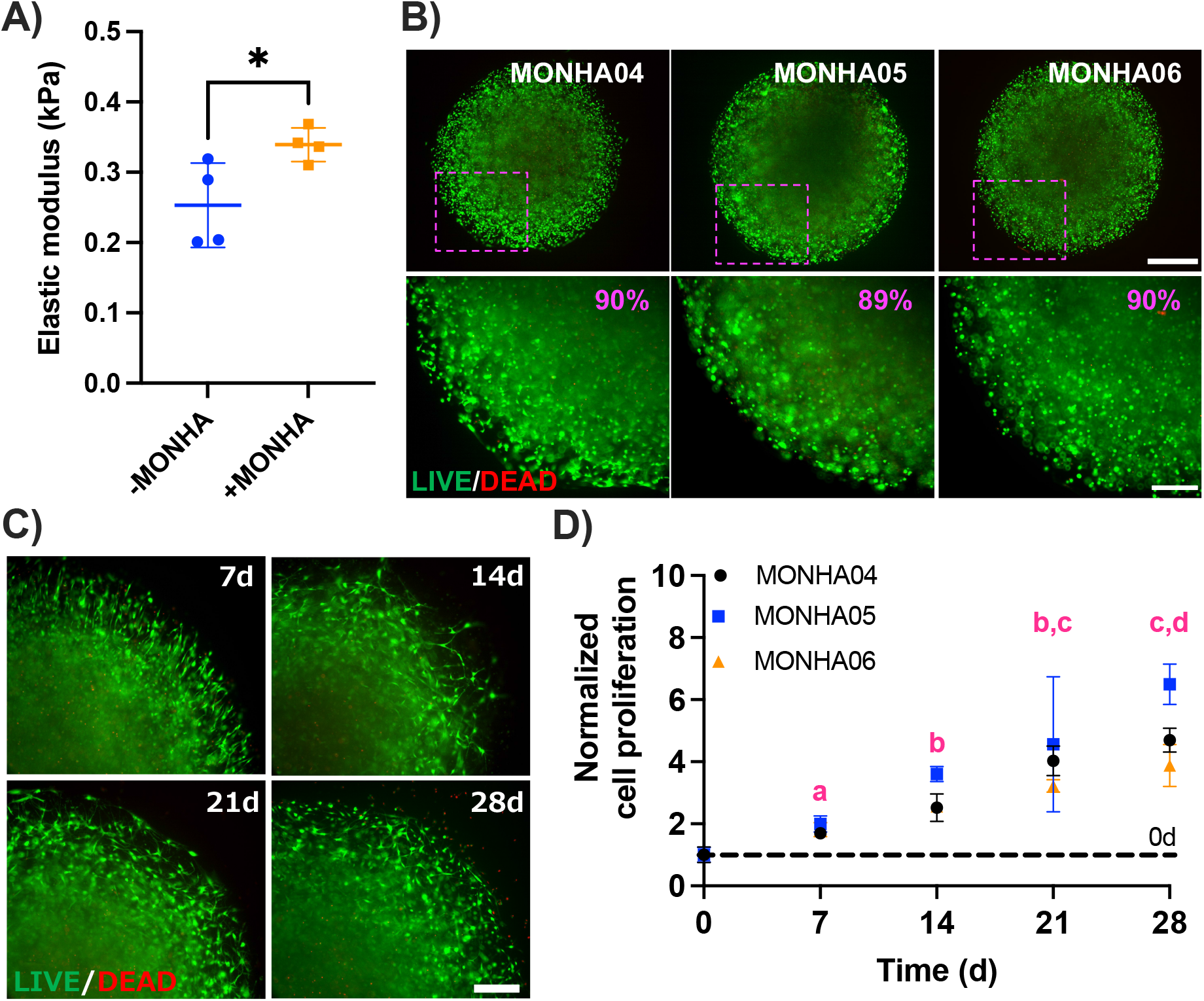
MONHA-encapsulated hydrogel stiffness, viability, and proliferation. (A) Elastic modulus of acellular and MONHA-encapsulated hydrogels (N = 4/group, *p < 0.05). (B) Live (green)/Dead (red) fluorescence images of 3 individual preparations (N = 3) at 0 d. Scale bars: 500 μm (top), 250 μm (bottom). (C) Longitudinal Live (green)/Dead (red) fluorescence images of representative hydrogels (MONHA05). Scale bar 250 μm. (D) Normalized cell proliferation over time (7 d, 14 d, 21 d, and 28 d; shared significance indicator letters represent non-significant difference (p > 0.05), distinct letters represent significant difference (p < 0.05)). Significance was determined by unpaired t-test (A) and two-way ANOVA using multiple comparisons tests (D) (*p < 0.05).

### 3.2 Astrocytes retain cell viability and proliferation over time

Immediately after UV crosslinking of MOHNA-encapsulated hydrogels (2.5 × 10^6^ cells/ml), we measured astrocyte viability. Live/Dead staining for 3 cell preparations (MONHA 04-06) demonstrated over 89% cell viability across strains (Fig. 2B). Cell viability was maintained over 4 weeks (Fig. 2C) and cells continued to significantly proliferate in a near linear fashion during that time (R^2^ = 0.94, 0.83, and 0.93 respectively) (Fig. 2D). We additionally encapsulated 5 × 10^6^ cells/ml within our hydrogels and similarly found high cell viability immediately after crosslinking and continued proliferation over 4 weeks in culture (Suppl. Fig. 2A-C).

### 3.3 Astrocytes develop typical stellate morphology over time in the hydrogel system

Astrocyte morphology is typically stellate in nature with small cell bodies and radial primary processes and branches connecting to other astrocytes via gap junctions (Sun et al., 2017; Cooper et al., 2018; Oberheim et al., 2009,). Astrocytes in our hydrogel system qualitatively demonstrated stellate morphology and extended processes and branches over time to promote interaction with neighboring astrocytes (Fig. 3A). Furthermore, after 4 weeks in culture MONHAs continued to express GFAP and CX43 (Fig. 3B), and thus, retained astrocyte-specific markers.

**Fig. 3.**
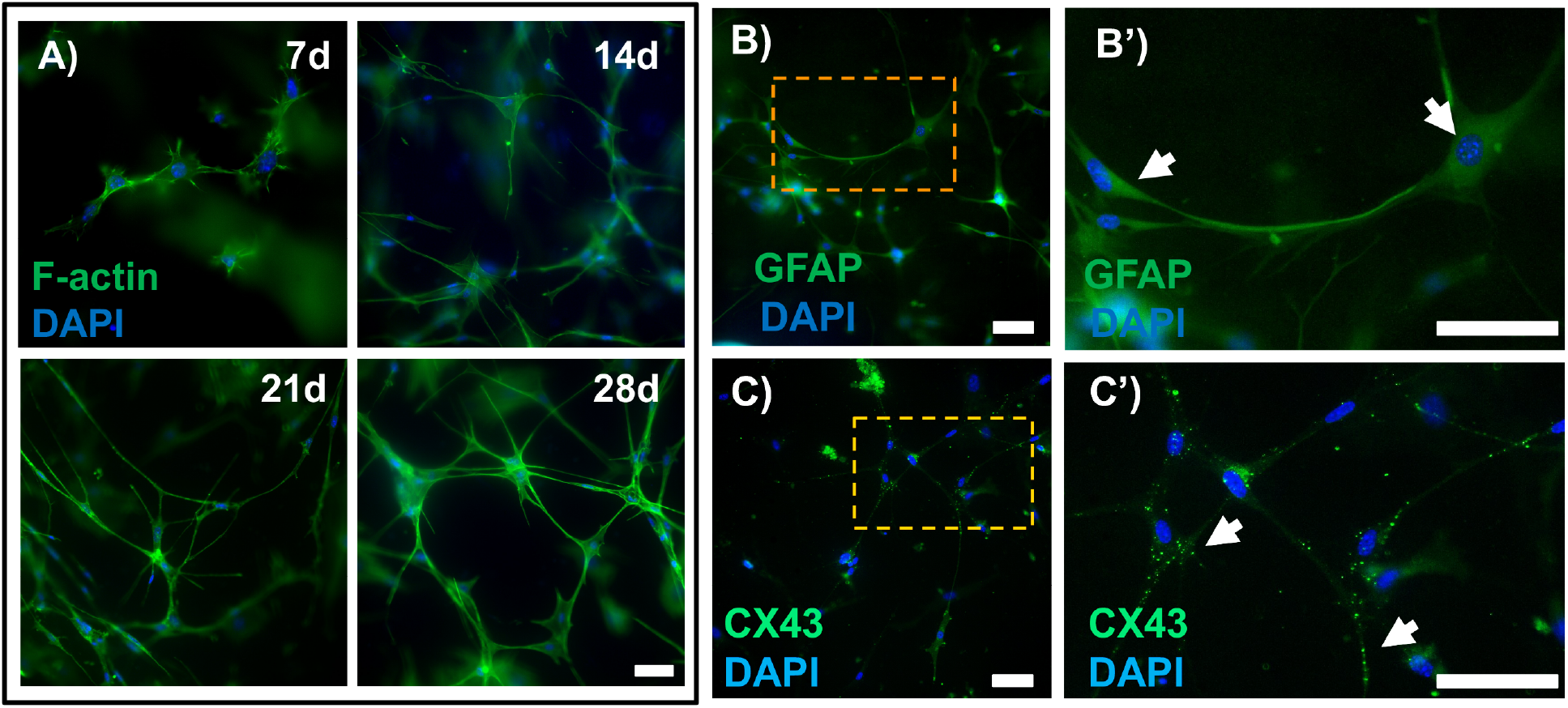
MONHA stellate morphology and astrocytic marker expression in hydrogels. (A) Representative fluorescence images of astrocyte morphology within hydrogels at 7 d, 14 d, 21 d, and 28 d. Scale bar: 100 μm. (B) Representative fluorescence images of astrocytes expressing GFAP (green) and (C) CX43 (green) in hydrogels at 28d. Scale bar: 100 μm. (B’-C’) Magnified images from boxed regions (orange) showing astrocytes expressing either GFAP (upper right, white arrows) or CX43 puncta (lower right, white arrows). Scale bar: 100 μm.

Confocal imaging of F-actin staining of individual cells within MONHA-encapsulated hydrogels were analyzed and illustrated process elongation over the course of three weeks (Fig. 4A-C). At 4 weeks, the rate of cell proliferation precluded analysis of distinct cells. Sholl analysis confirmed that increased time in culture significantly increased the length of astrocyte processes from 22.89 μm at week 1, to 52.65 μm at week 2 and 87.08 μm at week 3 (p < 0.0001) (Fig. 4D). In contrast, the degree of branching reflective of the number of total primary processes divided by the number of total branches per cell, remained unchanged over time, which is consistent with reports of cortical astrocyte-encapsulated hydrogels (Fig. 4E) (Placone et al., 2015; Butt et al., 1994). The two-way ANOVA analysis revealed significant main and interaction effects for the length of culture time and number of process intersections (i.e., complexity) away from center of cell body. In general, the effect of increased time in culture for 14 d and 21 d MONHA-encapsulated hydrogels was most prominently significant (p < 0.0001), with number of process intersections away from center of cell body demonstrating increased complexity as well (p < 0.05), in comparison to 7 d MONHA-encapsulated hydrogels. Therefore, systematic analysis of MONHA process complexity revealed enhanced complexity after 2-3 weeks in culture as compared to week 1 following encapsulation (Fig. 4F).

**Fig. 4.**
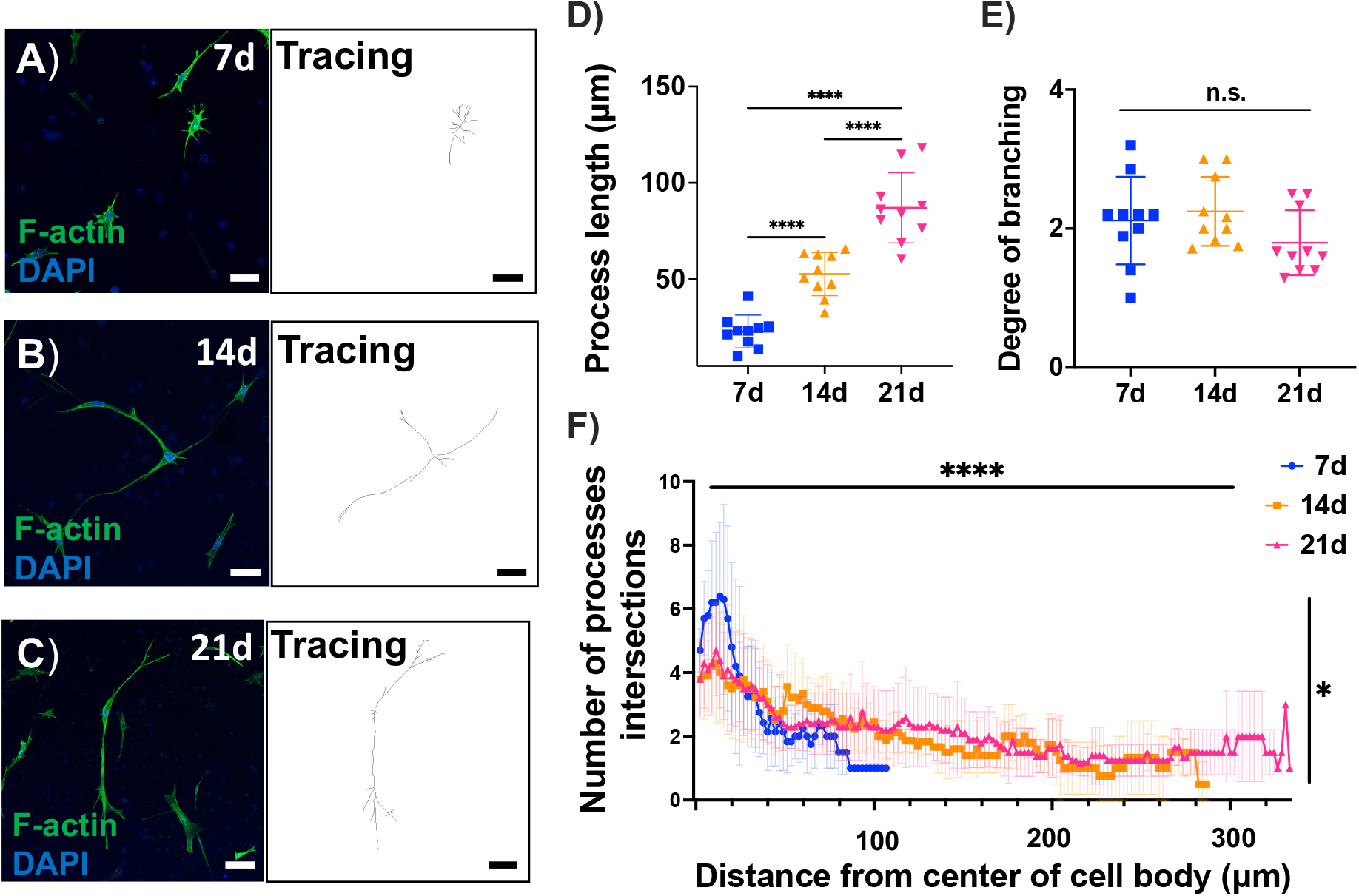
MONHA process length and branching in hydrogels. (A-C) F-actin staining and tracing of astrocytic morphology over time. Scale bar: 100 μm (A-B), 50 μm (C). (D-E) Process length and degree of branching of astrocytes. (F) Sholl analysis indicating the number of process intersections at each increasing radius from cell body for 7d, 14d, and 21d (N = 10 cells/group). Statistical significance was determined using one-way ANOVA (****p < 0.0001) for process length (D) and degree of branching (E), and two-way ANOVA main effects only model for process complexity over time (F) (main effect of time F (149, 2012) = 8.778, ****p < 0.0001, and main effect of process complexity F (2, 2012) = 3.992, *p < 0.05).

Taken together, these data support that 2-3 weeks culture within a 3D ECM matrix facilitates development of increased MONHA process length/complexity. MONHA-encapsulated hydrogels cultured for 3 weeks were used in all subsequent experiments.

### 3.4 TGFβ2 induces GFAP expression, actin cytoskeletal rearrangement, and ECM deposition in MONHA-encapsulated hydrogels

In glaucoma, astrocytes within the ONH become reactive and undergo remodeling of their F-actin cytoskeleton (Sun et al., 2017; Tehrani et al., 2016). Elevated TGFβ2 levels have been demonstrated within the glaucomatous ONH; likewise, TGFβ2 induces astrocyte reactivity in 2D culture (Hernandez et al., 2000; Pena et al., 1999; Prendes et al., 2013). Therefore, we asked whether exogenous TGFβ2 would induce astrocyte reactivity in our hydrogel system; to do so, we analyzed F-actin cytoskeletal levels, ECM protein deposition, and GFAP immunoreactivity.

MONHA-encapsulated hydrogels were cultured for 3 weeks prior to TGFβ2 treatment for 7 days (Suppl. Fig. 3A). Astrocytes retained viability and continued to proliferate across all groups (Suppl. Fig. 3B and C). MONHA-encapsulated hydrogels treated with TGFβ2 showed significant remodeling of F-actin networks (Fig. 5A) in a dose-dependent manner; an up to ∼ 6-fold increase in F-actin signal intensity compared to vehicle control-treated MONHA-encapsulated hydrogels was observed (Fig. 5B). Given the robust increase in F-actin intensity, treatment with 5 ng/ml TGFβ2 for 7 days was used for all subsequent experiments.

**Fig. 5.**
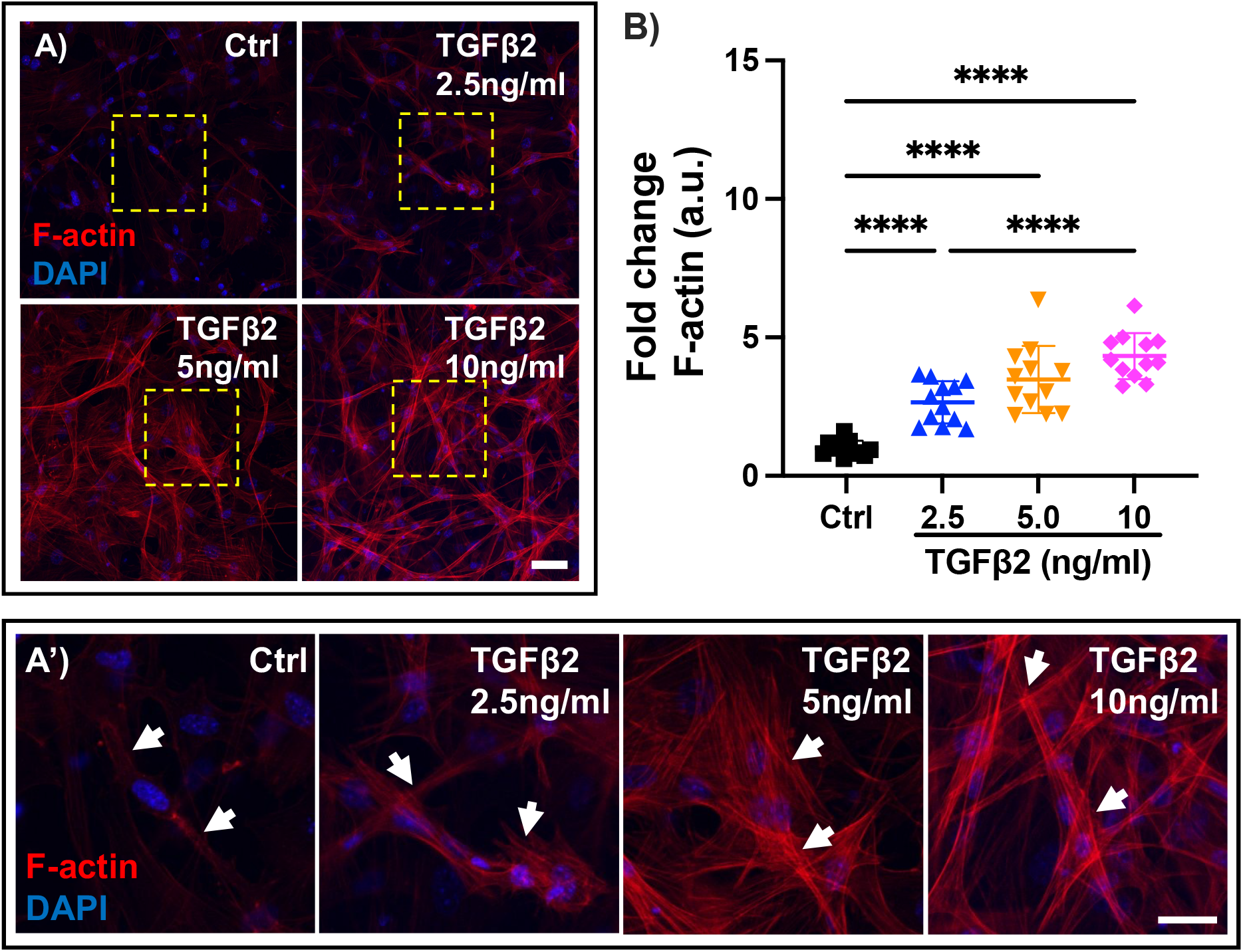
TGFβ2 effect on F-actin network in MONHA-encapsulated hydrogels. (A) Representative confocal fluorescence images of F-actin expression levels in control versus TGFβ2-treated MONHA-encapsulated hydrogels (2.5 ng/ml, 5 ng/ml, 10 ng/ml). Scale bar: 100 μm. (A’) Magnified images from boxed regions (yellow) of F-actin cytoskeletal changes (white arrows) in control versus TGFβ2-treated MONHA-encapsulated hydrogels (2.5 ng/ml, 5 ng/ml, 10 ng/ml). Scale bar: 50 μm. (B) Quantification of fold change in F-actin intensity (N = 10 fields of view/group). Statistical significance was determined using one-way ANOVA (****p < 0.0001) for F-actin expression levels (B).

;Reactive astrocytes upregulate ECM proteins and intermediate filament protein GFAP (Hernandez et al., 2000), and elevated TGFβ2 levels are associated with ECM remodeling within the glaucomatous ONH (Kim et al., 2017; Zode et al., 2011). We previously showed that MONHAs in conventional 2D culture increased GFAP expression and ECM deposition in response to TGFβ2 treatment (Kirschner, et al., 2021). Therefore, we asked whether exposure of MONHA-encapsulated constructs to 5 ng/ml TGFβ2 for 7 days would induce similar cellular responses in 3D culture.

Vehicle control treated MONHA encapsulated hydrogels showed low baseline levels of collagen IV and fibronectin deposition, and GFAP expression (Fig. 6A, B, C). In contrast, TGFβ2-treatment induced a significant ∼ 2.5-fold increase in collagen IV (p < 0.0001) and fibronectin (p < 0.001) signal (Fig. 6D and E). GFAP immunoreactivity increased ∼7.4-fold compared to controls (p < 0.0001) (Fig. 6F). Taken together, these data indicate that MONHAs encapsulated within our ECM hydrogel respond reliably to biochemical cues of glaucomatous insult (i.e., exogenous TGFβ2).

**Fig. 6.**
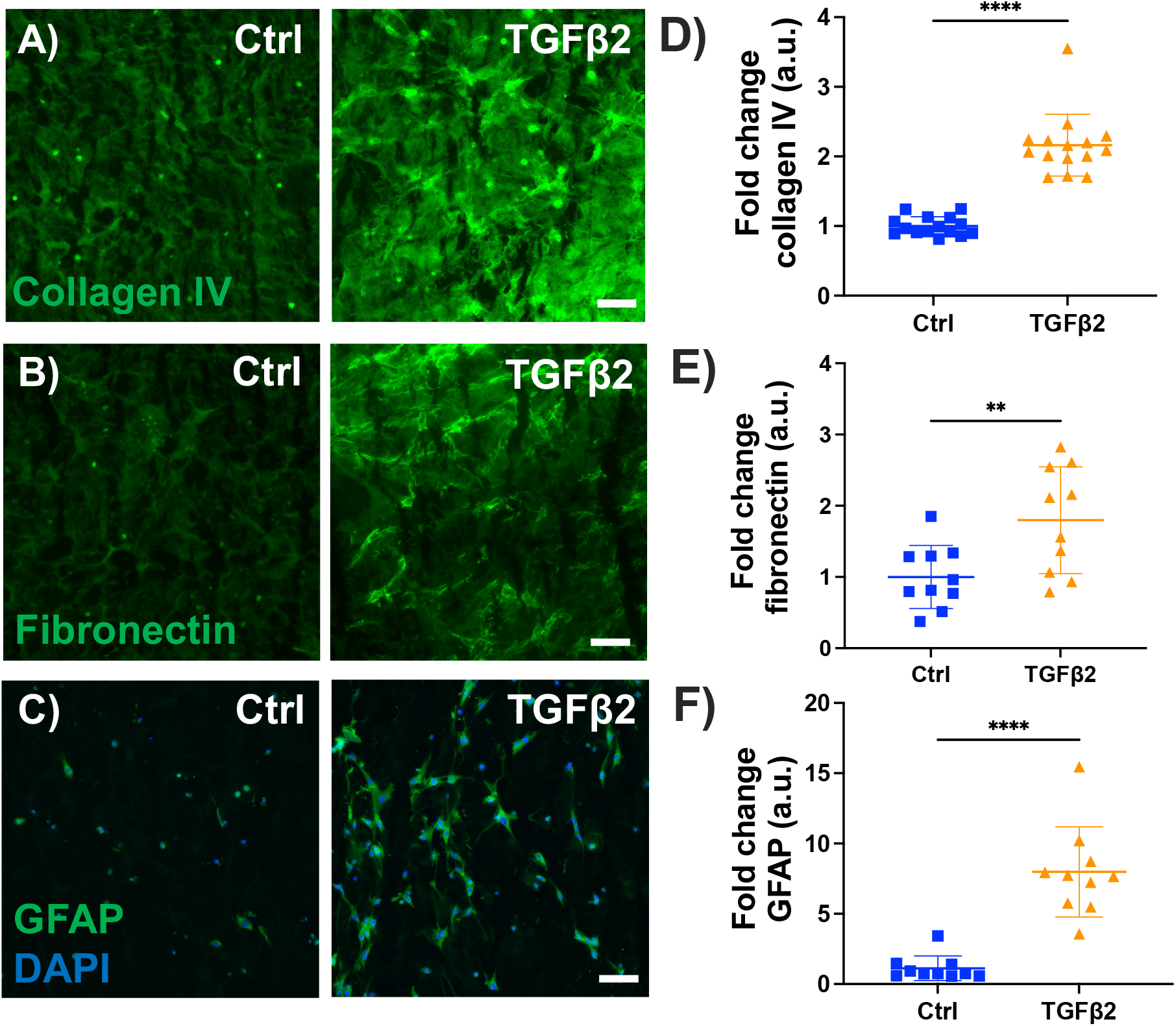
Effect of TGFβ2 on ECM protein and GFAP levels. (A-C) Representative fluorescence images of collagen IV, fibronectin, GFAP expression in control versus 5 ng/ml TGFβ2-treated MONHA-encapsulated hydrogels. Scale bar: 100 μm. (D-F) Quantification of fold change in signal intensity for collagen IV, fibronectin deposition, and GFAP expression shows significant difference between groups. (N = 5-7 fields of view/group for 2 strains). Statistical significance was determined using unpaired t-test (**p < 0.001, ****p < 0.0001).

## 4. Discussion

In glaucoma, elevated IOP progressively injures RGC axons within the ONH, leading to irreversible blindness. Evidence suggests that astrocytes residing within the ONH are one of the first responders to glaucomatous insult (Hernandez, 2000; Sun et al., 2017; Cooper et al., 2020; Wang et al, 2017). Initially, astrocyte reactivity provides support to RGCs via increased gap junction coupling and nutrient transfer (Sun et al., 2017; Blanco-Suarez et al., 2017; Cooper et al., 2020). However, there is evidence that excessive reactivity can become neurotoxic later in the disease and adversely affect the health of RGCs axons, partially by increasing ECM protein deposition and altering the astrocyte microenvironment (Clarke and Barres, 2013; Liddelow et al., 2017; Sterling et al., 2020). Animal studies have long investigated the dichotic astrocytic behavior *in vivo* providing crucial insight into astrocyte reactivity at the ONH, but much of the mechanistic details of the astrocyte response to glaucomatous insult remain unclear. In this study, our overarching goal was to engineer an *in vitro* system that allows for mechanistic analyses of astrocyte dichotic function in response to glaucomatous insult.

While many investigations on astrocyte behavior include conventional cell culture model systems, there are several limitations to traditional 2D culture, namely supraphysiologic substrate stiffnesses and the lack of a 3D scaffold. These limitations can translate into a less faithful representation of *in vivo* astrocyte star-shaped morphology and function. Many cells are intrinsically sensitive to substrate stiffness. For example, mesenchymal stem cells seeded on stiff substrates (>10 kPa) and conventional tissue culture plastic dishes retain mechanical information and behave differently to cells cultured on substrates similar in stiffness to human tissue (< 5kPa) (Yang et al., 2014; Heo et al., 2015; Price et al., 2021). As such, newer *in vitro* models seek to incorporate these nuances to more reliably model astrocyte behavior. Some groups have used viscoelastic polymer hydrogels to better model the 3D architecture of neural tissues. Since collagen and HA are rich within neural tissues, a majority of 3D hydrogels consist of collagen, collagen/HA, or collagen/HA/matrigel combinations to promote quiescent astrocyte stellate morphology (Placone et al., 2015). However, when specifically studying ONHA behavior, important caveats to these methodologies include both the slow polymerization rate (i.e., 15 – 30 minutes required for collagen/HA gel formation at 37 °C) and the batch-to-batch variability of matrigel (Caliari and Burdick, 2016). These aspects may affect ONHA viability and produce differences in the biochemical and mechanical properties of hydrogels from one preparation to another and thus, impact cellular behavior differently each time.

In order to circumvent hydrogel inconsistencies associated with slow polymerization rate and batch-to-batch polymer variability, we used short duration UV-mediated crosslinking and well-defined ECM proteins for hydrogel construction. We recently published on a UV-crosslinked trabecular meshwork hydrogel system using Irgacure as a photoinitiator, which allows rapid (seconds to minutes) crosslinking between photoactive ECM biopolymers (e.g., methacrylate-conjugated collagen type I, thiol-conjugated HA) (Li et al, 2021), and we sought to adapt this hydrogel for ONHA encapsulation. We formulated the hydrogel using a collagen:HA ratio of 3:1 for both its broad applicability across different cell types (Mazzocchi, Devarasetty et al. 2018, Mazzocchi, Devarasetty et al. 2019, Aleman, Sivakumar et al. 2021) and its recent association with reduced baseline cortical astrocyte reactivity (Placone et al., 2015). Incidentally, our initial studies using Irgacure yielded suboptimal MONHA viability, and thus, we incorporated the photoinitiator riboflavin (vitamin B2) within our system instead. Riboflavin is widely used for ocular collagen crosslinking for the treatment of keratectasia and can prevent excessive axial elongation in highly myopic eyes (Wollensak et al., 2003; Iseli et al., 2015). As a photoinitator, riboflavin may stabilize mechanical characteristics of collagen-based hydrogels while providing cytoprotective benefits (Ahearne and Coyle, 2016; Heo et al., 2015; Piluso et al., 2020). In our studies, use of 0.025% riboflavin and low UV intensity (10.3 mW/cm^2^) in the blue light range (405-500 nm) allowed for high ONHA viability after crosslinking for 5 minutes. Encapsulated MONHAs continued to proliferate over four weeks in culture. Moreover, the stiffness of this ECM hydrogel was well within the range of *in vivo* neural tissues (Budday et al., 2015, Budday et al., 2017), in stark contrast to widely used 2D culture systems of supraphysiologic stiffness.

ONH astrocytes *in vivo* are stellate in morphology with extended and branched processes to promote coupling with neighboring astrocytes and RGC axons for neurotrophic support (Sun et al., 2017; Cooper et al., 2018; Oberheim et al., 2009). To investigate whether MOHNAs would develop such a stellate morphology within our hydrogel system, we measured cell process elongation and complexity over time using Sholl analysis. We confirmed that MONHAs encapsulated within hydrogels possess typical star-shape morphology with increased processes extension and complexity over 28 days in culture. Moreover, MONHAs cultured for this duration expressed CX43 in processes branching to neighboring astrocytes. These findings support the use of this 3D hydrogel system to study process remodeling, gap junction communication, and nutrient transfer between astrocytes in response to glaucomatous insult.

Astrocyte reactivity and gliosis is a complex process that is not easily defined. Astrocyte reactivity encompasses a spectrum of activated states that include both neuroprotective and neurotoxic phenotypes (Liddelow et al., 2017; Escartin et al., 2021). In general, astrocyte reactivity is characterized by morphologic changes such as enlargement of soma/cell shape and cytoskeletal remodeling with thicker protrusive processes, as well as functional changes that alter the cell’s microenvironment (Molofsky et al., 2021; Vecino et al., 2016). In glaucoma, ONH astrocyte reactivity is associated with intrinsic F-actin remodeling, increased ECM protein deposition and upregulation of GFAP (Hernandez et al., 2000). To determine whether MONHA reactivity could be reliably induced in our system, we treated MONHA-encapsulated hydrogels with TGFβ2, a known glaucomatous stressor (Kim et al., 2017; Pena, et al. 1999; Zode et al., 2011). We observed that TGFβ2-treated MONHAs developed increased F-actin signal intensity, elevated fibronectin, and collagen IV protein deposition, and increased intracellular GFAP levels consistent with our previous work in 2D culture (Kirschner et al., 2021). Importantly, our hydrogel system enables – for the first time -accurate analyses of astrocyte morphological changes and the interplay between the surrounding ECM and astrocyte behavior in a relevant 3D microenvironment.

In conclusion, we have engineered a biomimetic MONHA-encapsulated hydrogel system to investigate key aspects of ONH astrocyte behavior in response to biochemical cues of glaucomatous insult that cannot be easily assessed via 2D cell culture. An additional advantage of the 3D MONHA-encapsulated hydrogel described herein is the ability to study astrocyte response to biophysical insults (i.e., matrix stiffening and biomechanical stress). Age-associated stiffening and compressive/tensile strains conferred by elevated IOP are thought to contribute significantly to MONHA reactivity in glaucoma (Sigal et al., 2007; Grytz et al., 2012; Korneva et al., 2020), but the mechanisms underlying this response are unknown. We recently described a role of mechanosensitive channel activation in TGFβ2-induced MONHA dysfunction in 2D culture (Kirschner et al., 2021). In future experiments, we aim to use this 3D culture system to study how mechanosensitive channel activity may modulate MONHA response to glaucomatous biophysical cues.

## Acknowledgments

We thank Dr. Alison Patteson at Syracuse University for rheometer access, Drs. Audrey M. Bernstein and Mariano S. Viapiano, and the Neuroscience Microscopy Core at Upstate Medical University for imaging support, and Mona El Gendi for assisting with cell characterization.

## Author contributions

A.N.S., A.K, H.Y., A.S., T.B., H.L., P.S.G., and S.H. designed all experiments, collected, analyzed, and interpreted the data. A.N.S and P.S.G. wrote the manuscript. All authors commented on and approved the final manuscript. P.S.G. and S.H. conceived and supervised the research.

## Data and materials availability

All data needed to evaluate the conclusions in the paper are present in the paper and/or the Supplementary Materials. Additional data related to this paper may be requested from the authors.

