## Supplemental Figures for "Engineering a 3D hydrogel system to study optic nerve head astrocyte morphology and behavior in response to glaucomatous insult"

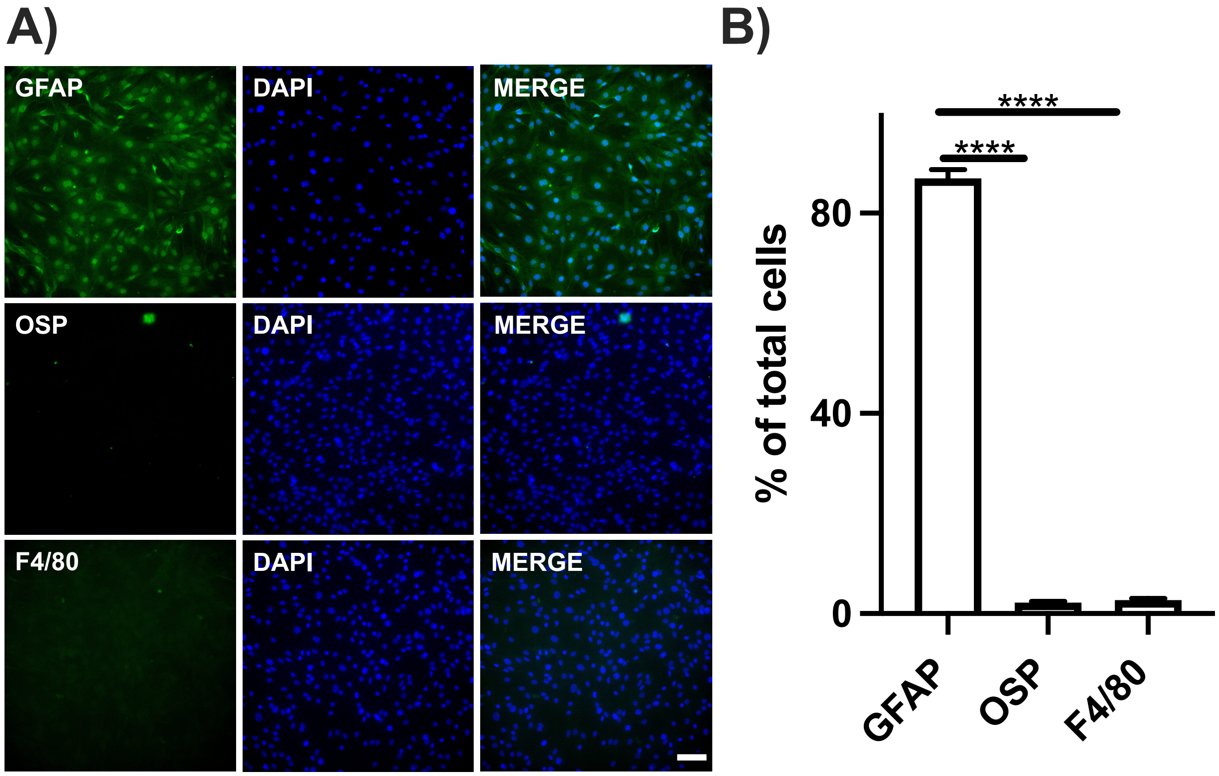


**Fig. S1. MONHA cell characterization.** (A) MONHAs were fixed and probed with antibodies against GFAP (green), OSP (green), and F4/80 (green). The cells were counterstained with DAPI to label DNA (blue) as a marker for nuclei. Scale bar: 100 μm. (B) Quantitative analysis shows that more than 85% of the cells in culture express GFAP. Statistical significance was determined using one-way ANOVA (**** $p < 0.0001$).

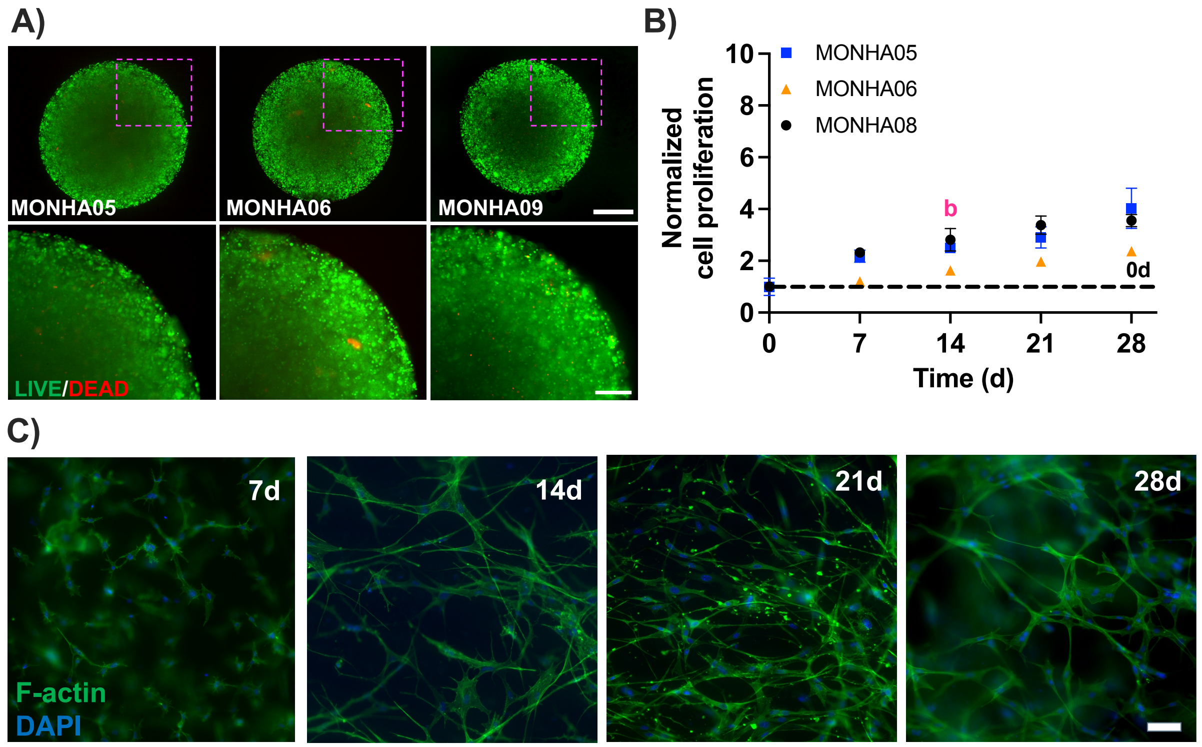


**Fig. S2. MONHA-encapsulated hydrogel stiffness, viability, and proliferation.** (A) Live (green)/Dead (red) fluorescence images of 3 individual preparations (N = 3, 5 x $10^6$ cells/ml hydrogel) at 0 d. Scale bars: 500 µm (top), 250 µm (bottom). (B) Normalized cell proliferation over time (7 d, 14 d, 21 d, and 28 d; shared significance indicator letters represent non-significant difference ($p > 0.05$), distinct letters represent significant difference ($p < 0.05$)). (C) Representative F-actin staining of astrocytic morphology over time (MONHA05 at 7 d, 14 d, 21 d, and 28 d). Scale bar: 100 µm.

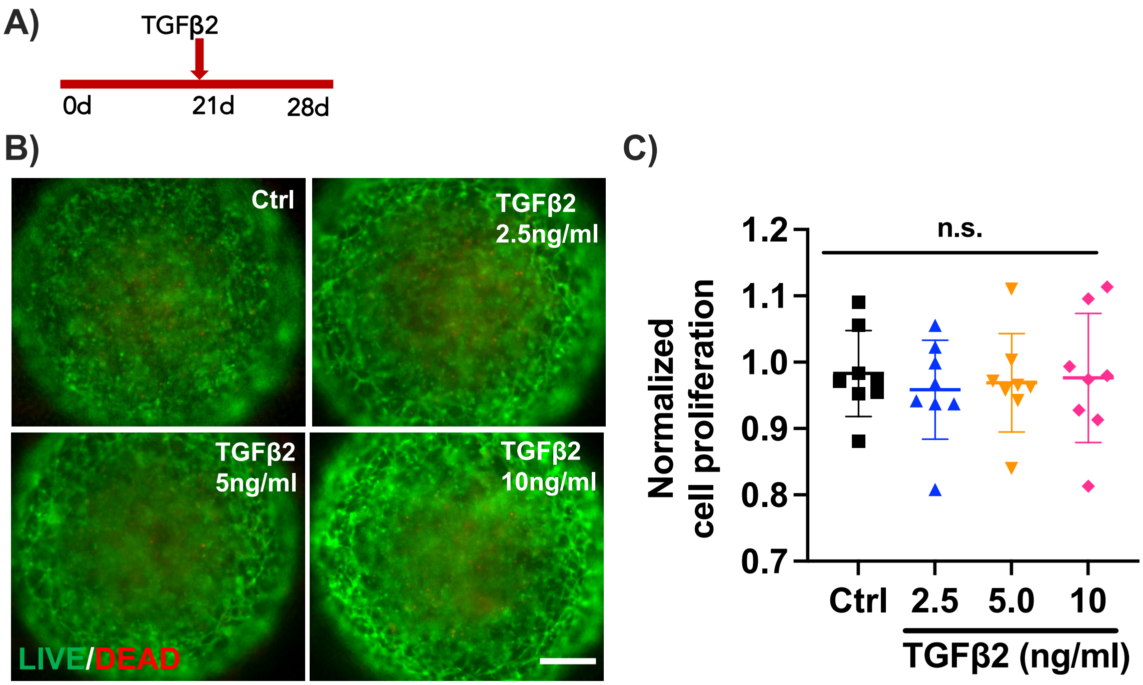


**Fig. S3. TGFβ2 effect on MONHA-encapsulated hydrogel viability and proliferation.** (A) Experimental timeline. (B) Representative fluorescence images of Live (green)/Dead (red) stain in control versus TGFβ2-treated MONHA-encapsulated hydrogels (2.5 ng/ml, 5 ng/ml, 10 ng/ml). Scale bar: 500 µm. (C) Normalized cell proliferation with increasing TGFβ2 concentrations (0, 2.5, 5.0, or 10 ng/ml, N = 4/group).
